# Gene inversion increases evolvability in bacteria

**DOI:** 10.1101/293571

**Authors:** Christopher Merrikh, Houra Merrikh

## Abstract

In bacteria, most genes are encoded on the leading strand, co-orienting the movement of the replication machinery with RNA polymerases. This co-orientation bias reduces the frequency of highly detrimental head-on collisions between the two machineries. This and other work set up the expectation that over evolutionary time, head-on alleles are selected against, maximizing genome co-orientation. Our findings challenge this model. Using the well-established GC skew method, we reveal the evolutionary inversion record of all chromosomally encoded genes in multiple divergent bacterial pathogens. We find that a surprisingly large number of co-oriented genes have inverted to, and are retained in the head-on orientation. Furthermore, we find that these head-on genes, (including key antibiotic resistance and virulence genes) have higher rates of nonsynonymous mutations and are more frequently under positive selection (dN/dS>1). Based on these results, we propose that bacteria increase their evolvability through gene inversion and promotion of head-on replication-transcription collisions.

## Introduction

DNA replication and transcription occur simultaneously, leading to collisions between the replisome and RNA polymerases (RNAPs)^1–3^. These conflicts collapse the replication fork multiple times per cell cycle in bacteria, breaking the DNA, and increasing mutagenesis^4–9^. Therefore, conflicts can significantly influence fitness, replication speed, and genome integrity. Cells harbor essential, conserved factors dedicated to resolving conflicts^9–14^. However, despite these mechanisms, recent work has shown that conflicts still occur frequently, precipitating the need for replication restart outside of the origin, *oriC*, multiple times per cell cycle^8,15^.

Replication-transcription conflicts can occur in either of two orientations. Co-directional (CD) encounters can occur within genes encoded on the leading strand when DNA replication forks overtake RNAPs^16^. The consequences of these encounters are far less severe than head-on (HO) conflicts, which occur in genes encoded on the lagging strand^3,4,13,17^. Notably, head-on conflicts cause an increase in local mutation rates. In keeping with the more detrimental nature of head-on conflicts, all known bacterial genomes display an orientation bias in which 54% to 82% of genes are co-oriented with replication. Among highly transcribed genes, the case is more exaggerated. In particular, rRNA operons are co-directionally oriented in all known bacterial species^1,17^. Previous analyses have also shown that essentiality is a major driver of co-orientation, suggesting that highly conserved genes are excluded from the lagging strand^18,19^. These findings implicate head-on replication-transcription collisions as a potent driver of genome organization. This mechanism is anticipated to reduce mutagenesis of both essential (essentiality as determined under laboratory conditions) and conserved (core) genes. In principle, the use of co-orientation as a means of protecting cells from head-on conflicts should apply to other genes as well.

It is unclear how the existing co-orientation bias of bacterial genomes arose. Therefore, important questions about the evolutionary history of genome architecture, and the impact of replication-transcription conflicts have yet to be answered. Generally speaking, three patterns could have led to the existing co-orientation bias: 1) A reduction in the number of head-on genes from a more balanced ancestral genome, 2) An increase in the number of head-on genes on an ancestral genome that was predominantly composed of co-directional genes, or 3) A static balance between head-on and co-directional genes that has been preserved over evolutionary time. Our understanding of conflicts predicts that the first model is more likely to be accurate because negative selection should generally reduce the abundance of head-on alleles/conflicts. Though these model address fundamental questions about evolution, they have not been tested.

The literature demonstrates that local DNA inversions can spontaneously occur in all living cells^20–33^. Previous work shows that such inversions can be identified by local changes in the GC skew, the well-established asymmetric distribution of guanine versus cytosine along single DNA strands^22,24,34^. The mutational footprint of normal DNA replication results in a positive G:C ratio, and a positive GC skew value within the leading strand of the replication fork (GC skew = (G-C)/(G+C))^35,36^. In essence, the GC skew of any given DNA region reveals its long term orientation with respect to the leading strand of the replication fork. As such, local inversions in the GC skew (regions in which cytosine outnumbers guanine) are a strong indication that a physical inversion has occurred^24,32^. Though some sequences may naturally defy the typical GC skew pattern over short lengths due to functional constraints, this pattern is generally robust. Therefore, gene inversion models are testable.

To our knowledge, the GC skew has never been used to systematically identify gene inversions genome-wide. Here, we analyze the GC skew of all chromosomally encoded genes to reveal the global gene inversion record of multiple important clinical pathogens. Our data demonstrate that, contrary to expectations, bacterial genomes are consistently gaining new head-on genes/operons via inversion. This likely increases the frequency of head-on replication-transcription conflicts, which are known to increase gene-specific mutation rates. Using comparative evolutionary analyses, we show that head-on genes have elevated rate of retained nonsynonymous mutations, compared to co-directionally oriented genes across bacterial phyla. Furthermore, we find that positive selection acts on a higher percentage of head-on genes, suggesting that the head-on orientation can actually be beneficial. Based on these findings, we propose that bacteria are universally increasing the mutability of their genomes via co-directional to head-on gene inversions. In particular, we find that many virulence and antibiotic resistance genes recently inverted to, or have long been retained in the head-on orientation. This suggests that pathogens commonly harness the mutagenic capacity of head-on conflicts to accelerate the evolution of hyper-virulence and antibiotic resistance.

## Results

### Physical DNA inversions result in an inverse GC skew

As previously shown in a variety of bacterial species, guanine nucleotides outnumber cytosine nucleotides on the leading strand of each arm of the chromosome. Using an example sequence, we demonstrate that the results of the GC skew calculation are inverted as the direct consequences of a physical DNA fragment flip (Fig. 1A). For all known species, the GC skew is, on average, positive with respect to the leading strand of DNA replication forks. In Figure 1B, we show the *M. tuberculosis* GC skew as an example (GC skew values are indicated in green/purple). Despite the generally positive GC skew values on each arm, large regions with inverse GC skew values are clearly visible as deviations from the strand averages (Fig. 1B, gray arrows). The GC skew map demonstrates that numerous local inversions that have occurred over the course of natural evolution in *M. tuberculosis* and potentially other species. To further assess the validity of this method of inversion detection, we compared 55 fully assembled *M. tuberculosis* genomes to identify chromosomal fragments that have inverted relative to the H37Rv reference strain. We then compared the GC skew of single gene regions before and after inversion (Figure 1C). These data clearly demonstrate that physical DNA fragment inversion concomitantly inverts the GC skew, irrespective of original or final orientation of the region (CD to HO, or, HO to CD). As such, negative GC skew values (with respect to the leading strand) are a direct indication of gene inversion.

**Figure 1.**
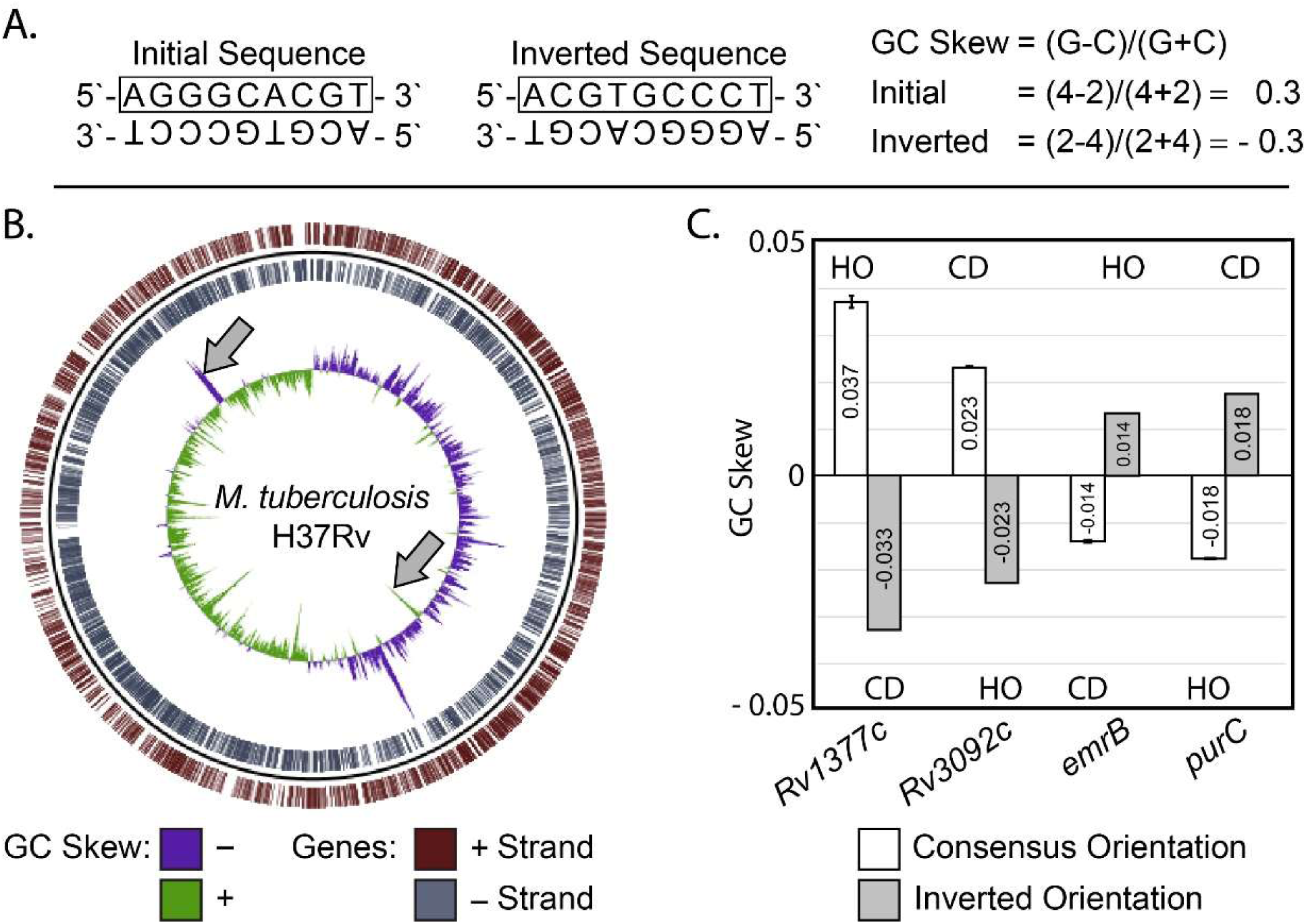
DNA sequence inversion concomitantly inverts GC skew values. A) The effect of DNA fragment inversion on GC Skew. An example DNA sequence and its inverted form are depicted. B) GC skew values along the top strand are plotted on the *M. tuberculosis* H37Rv genome. Positive GC skew values are plotted in purple, and negative values are in green. Potential inversions are apparent as deviations from the local average (examples are indicated by gray arrows). The box indicates the the upper strand is used for GC skew calculation by convention. The GC calculation on the right shows that the relative abundance of guanine and cytosine residues in the top strand flips upon DNA inversion, resulting in an inverse GC skew value. C) Anecdotal examples of four genes that naturally inverted in 1 out of 55 analyzed genomes. Here we assessed the orientation of all genes in fully assembled genomes by comparing the coding strand of each gene relative to the positions of *ori* and *ter* for each genome. The orientation of each allele with respect to the movement of the DNA replication fork is indicated above/below their corresponding data columns. Head-On alleles (encoded on the lagging DNA strand) are marked “HO”, and co-directional alleles (encoded on the leading DNA strand) are marked “CD”. Columns in white represent the consensus orientation values identified in 54 of the 55 analyzed *M. tuberculosis* genomes. Columns in gray represent an inverted allele identified in a single lineage. GC values or averages are marked on the corresponding columns.

### Representative genomic regions suggest that inverted GC skews correlate with genes in the head-on orientation

As an initial assessment of bacterial DNA inversion patterns, we calculated the chromosomal GC skew of multiple divergent bacterial species (see complete list of species and name abbreviations in Table S1). We then conducted a preliminary visual investigation of various chromosomal regions in which negative GC values predominate (Fig. 2, gray). We immediately noticed that these regions tend to correlate with the presence of head-on genes or operons. This suggests that many head-on genes and operons may have originated via the inversion of co-oriented alleles. To better assess this possibility, we re-calculated the average GC skew over whole genes rather than analyzing arbitrary step sizes on the chromosome (Fig. 2, black boxes). This reduces noise in the GC skew signal and reveals the average GC skew values of single genes. These data support our initial impression that head-on genes frequently have negative GC skew values, whereas most (though certainly not all) co-oriented genes have positive GC skew values.

**Figure 2.**
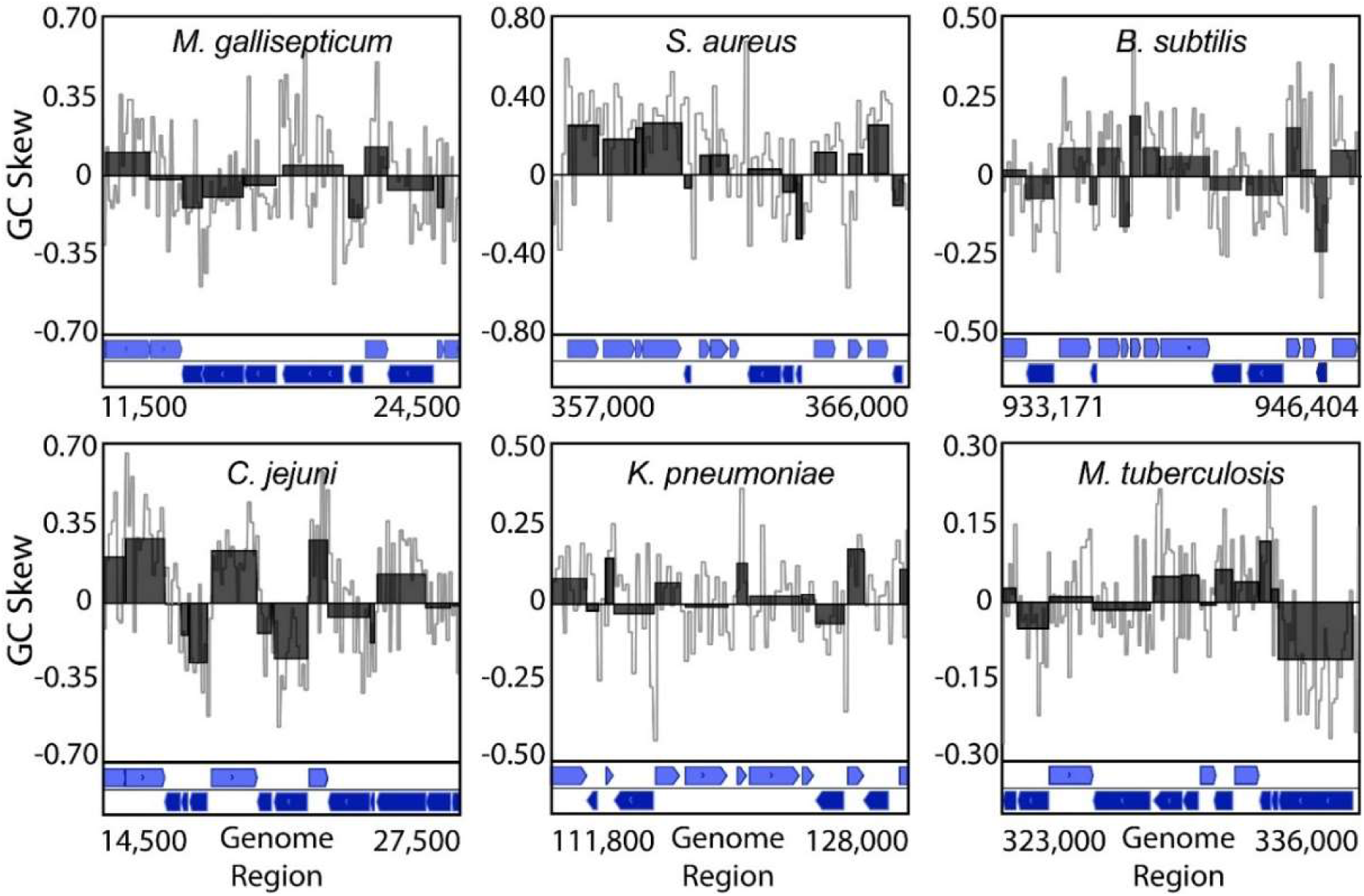
Representative genomic regions show that inverted GC skews correlate with genes in the head-on orientation. High resolution (100 bp step size) GC skew values are shown at in gray, or as an average over whole genes (black). The locations of individual genes and their coding strand are indicated below. For each species, head-on genes are annotated on the lower box (transcribed right to left, dark blue), and co-directional genes are on the upper box (light blue).

### A major percentage of head-on genes were originally co-directional

To further probe the correlation between head-on genes and negative GC skew values, we used a genome-wide quantitative analysis. Here, we binned GC skew data for each gene by orientation, and again by value: either >0 or ≤0 (Fig. 3). This second cutoff should reveal the proportion of either co-directional or head-on genes that have been retained in their current orientation versus those that are the product of an inversion event. For all species, these data are consistent with our initial qualitative observations: Many head-on genes have a negative GC skew, indicative of recent inversion from the opposite orientation. In most of the species, this applied to the majority of head-on genes. Conversely, few co-directional genes have a negative GC skew. These data demonstrate that a significant percentage of head-on genes are the product of inversion. This directly opposes the expectation that the production of head-on genes should be rare due to their ability to cause harmful head-on replication-transcription conflicts.

**Figure 3.**
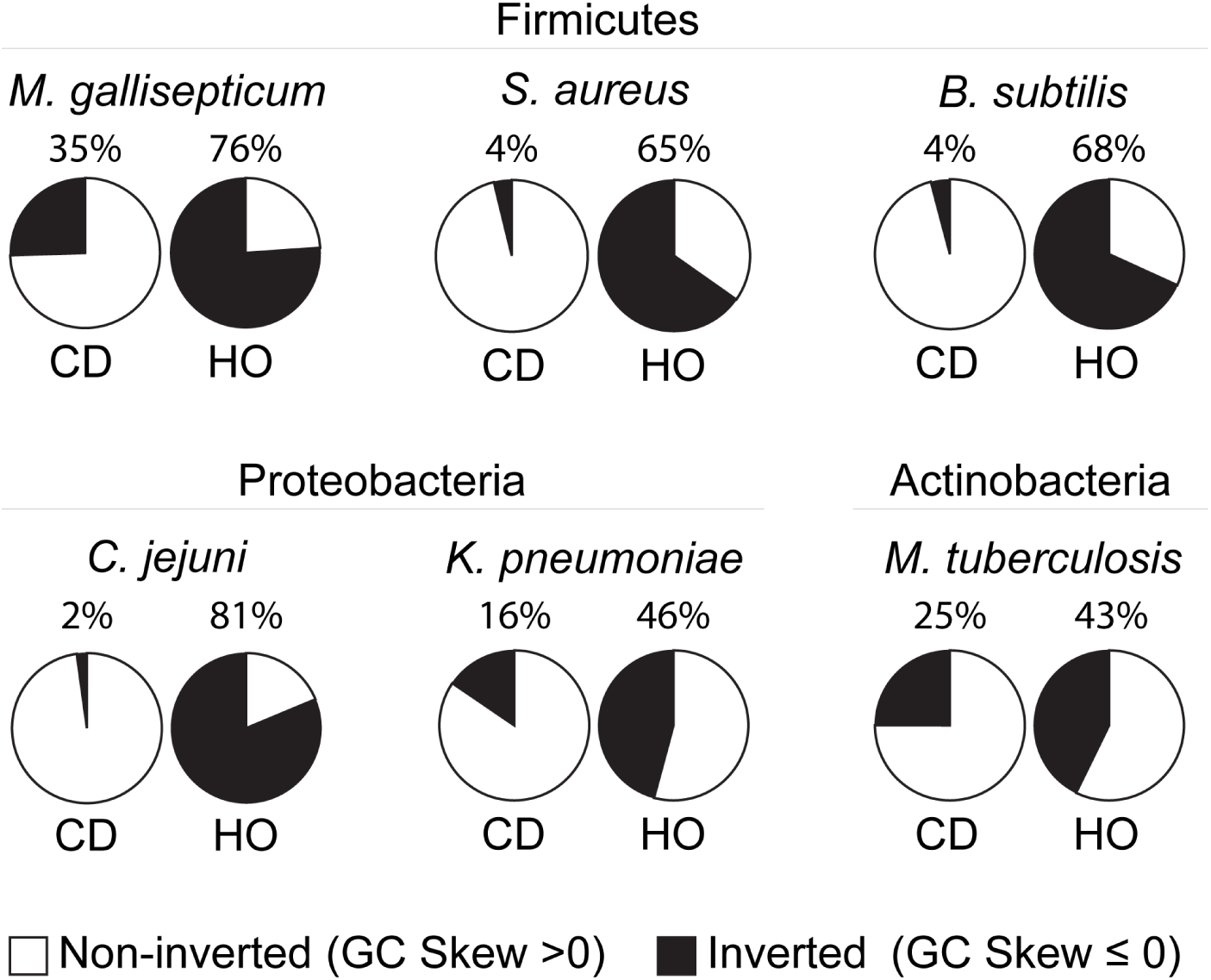
A large percentage of head-on genes were originally co-directional. The GC skew values of individual genes were binned by current orientation, and again by GC skew value. Positive GC Skew values indicate long term retention in the current orientation. Negative GC skew values indicate that a gene has recently inverted from the opposite strand/orientation. Leading strand genes are labelled “CD” as they are co-directionally oriented with the DNA replication fork, whereas lagging strand genes are labelled “HO” as they are head-on to DNA replication. GC skew values are also presented in the form of column graphs in Figure 5.

**Figure 5.**
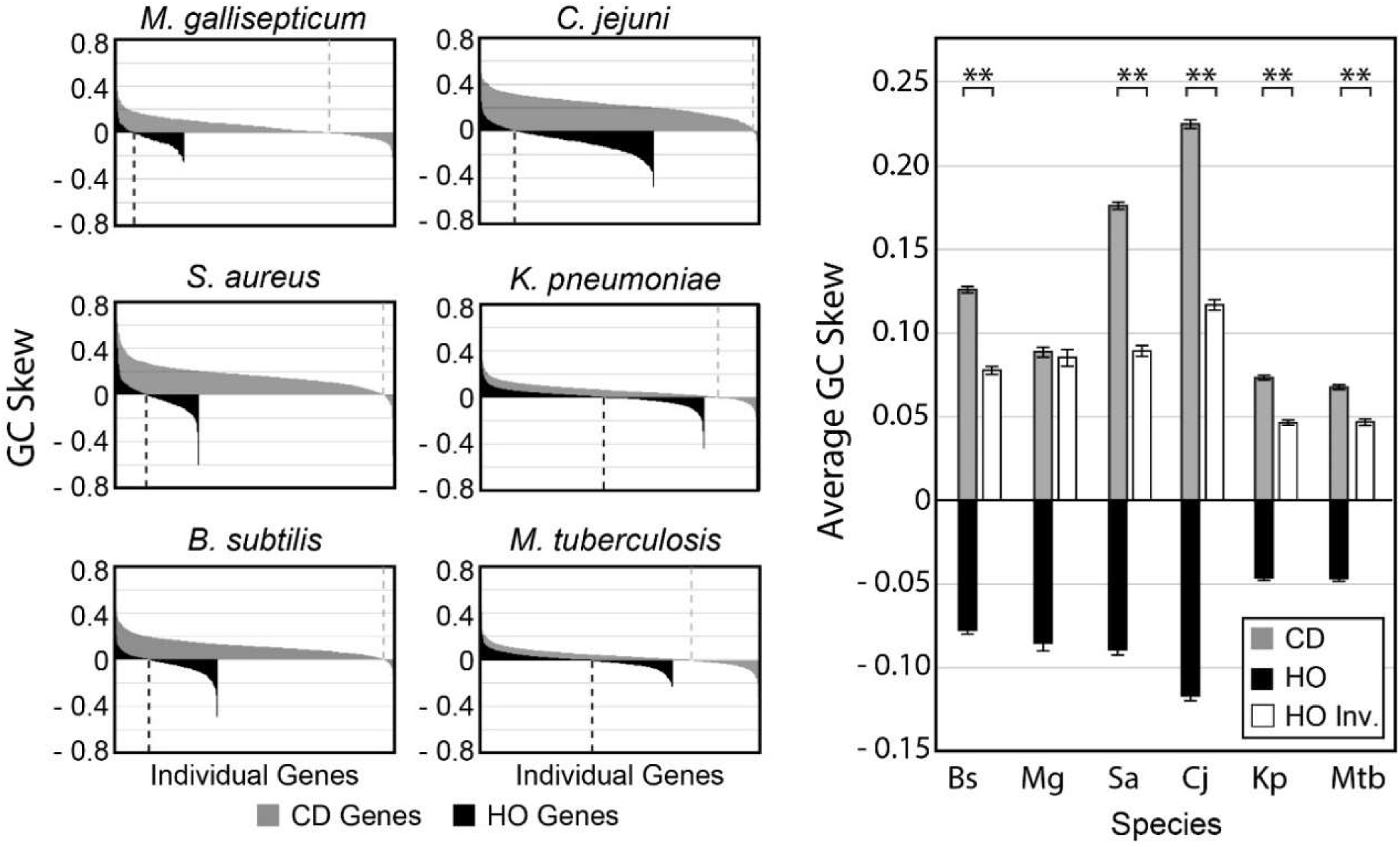
Inverted head-on genes are retained following inversion. Left) The distribution of GC skew values for all co-directional (leading strand, gray) or head-on (lagging strand, black) genes are plotted, one column per gene. Columns are sorted from highest to lowest GC value resulting in the appearance of a curve. Values below zero represent inverted genes. Values above zero suggest long-term orientation conservation. Dashed lines highlight the inflection point between positive and negative value in each group. Right) The magnitude of the average GC skew of inverted head-on genes is diminished relative to retained co-directional genes. The average GC skew of co-directional genes with a positive GC skew value (gray) or head-on genes with a negative GC skew value (black) are plotted. For side-by-side comparison of the magnitude of the head-on and co-directional averages, the head-on values were manually re-plotted as positive values (white). Error bars represent the standard error of the mean. Significance was determined using the Z-test. ** indicates p < 0.0001.

### Genes in the head-on orientation have a higher nonsynonymous mutation rate

The wide-spread and frequent creation of new head-on genes is somewhat counterintuitive as it should create new head-on replication-transcription conflicts. Presumably, cells harboring these new head-on alleles should experience increased replication stress and divide more slowly, resulting in negative selection pressure. Potential explanations for these contradictory findings include the possibility that the new head-on genes are not expressed during replication, thereby avoiding conflicts. It is also possible that the head-on orientation confers some benefit outweighing the detrimental effects of conflicts. Given that many of these genes are highly conserved, and at least some are definitively expressed during replication, it is likely that there is a benefit to having certain genes in the head-on orientation. Consistent with the latter possibility, we previously showed that head-on conflicts increase gene specific mutation rates and accelerate evolution in *B. subtilis* in nature (REF). Based on these observations, we proposed that head-on conflicts in certain genes may provide a net benefit to the cells through the more rapid creation of advantageous mutations, overcoming the negative selection pressure caused by the conflict. However, it is unclear whether accelerated head-on gene evolution occurs outside of *B. subtilis.*

To assess the mutation rates of head-on and co-directional genes, we performed genome-wide mutational analyses for each species *in silico* using TimeZone software (Fig. 4). Specifically, we analyzed the rates of non-synonymous (dN) and synonymous (dS) mutations among the core genes of each species, using at least 10 whole genomes for each. Our analysis shows that the rate of non-synonymous mutations is elevated in the head-on genes of each species relative to the corresponding co-directional genes (Fig. 4). Likewise, the average dN/dS ratio is also significantly elevated for head-on genes. These data are consistent with our initial analysis in *B. subtilis*, and establish a broadly conserved pattern across three bacterial phyla in which head-on genes are evolving at an accelerated rate in nature. Therefore, the ongoing co-directional to head-on inversions observed across bacterial species should increase the average mutation rate for many genes in each species.

**Figure 4.**
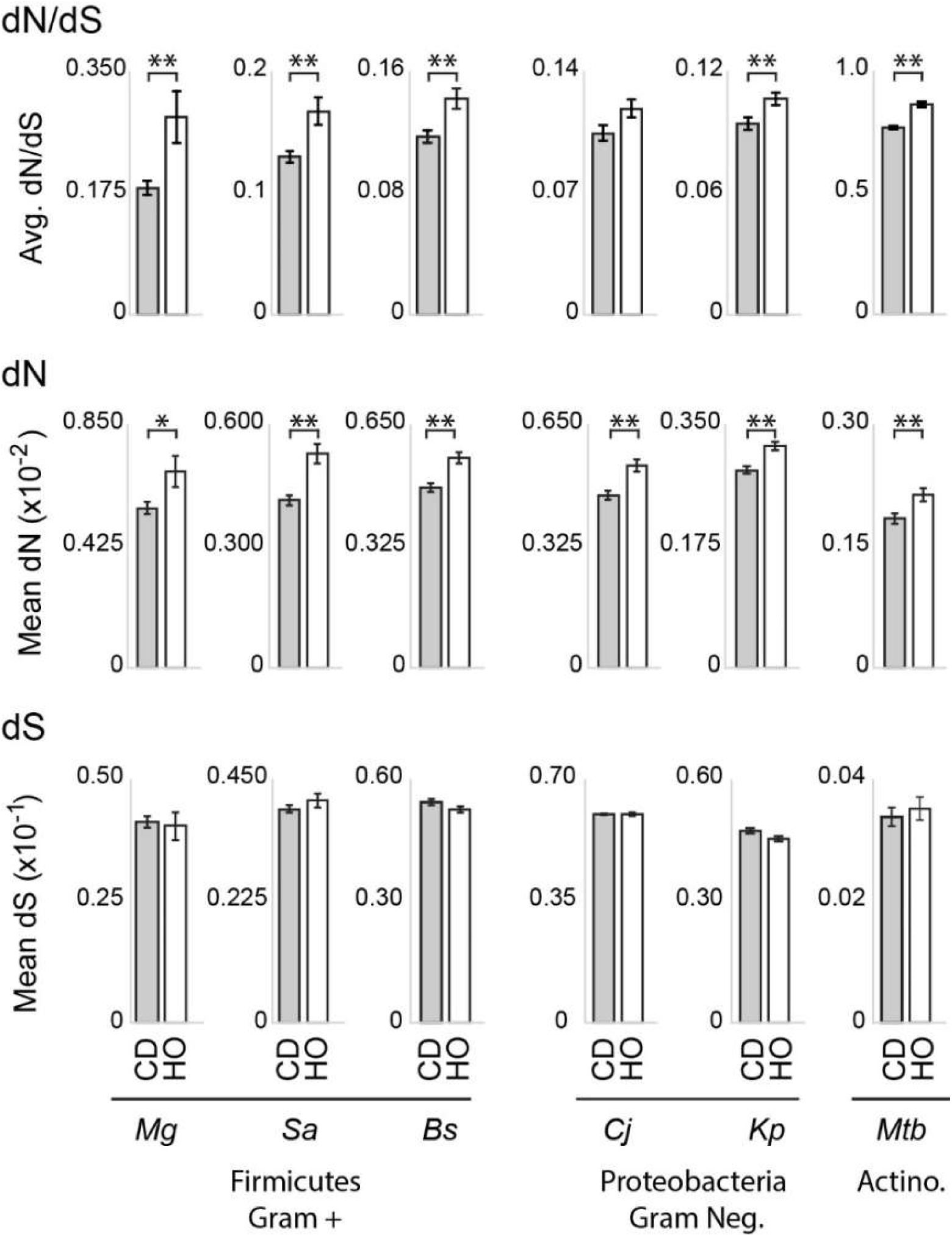
Head-on genes evolve at an accelerated rate across bacterial phyla. Whole genome mutational analysis including the dN/dS ratio, rate of nonsynonymous mutations (dN), and rate of synonymous mutations (dS) are shown. Values were calculated for core genes that are ≥ 95% conserved in length and amino acid sequence using at least 10 complete genomes per species. Additional species information and abbreviations are explained in Table S1. Error bars represent the standard error of the mean. Statistical significance was determined by the two-tailed Z-test. * indicates *p* < 0.05. ** indicates p < 0.01.

### Positive selection acts on a higher percentage of head-on genes

Though we find that head-on genes have a higher mutation rate, mutations are not necessarily beneficial. In other words, the elevated dN and dN/dS ratio is consistent with increased positive selection which would support the beneficial hypothesis, and decreased negative selection acting on head-on genes which would not. To better understand the relative influence of different selective pressures, we searched for overt signs of positive selection. Specifically, we calculated the frequency of head-on or co-directional genes for which a dN/dS ratio of greater than 1 was observed. This high ratio of nonsynonymous to synonymous mutations should only occur if a gene is experiencing strong positive selection. We initially observed a higher frequency of genes with dN/dS values exceeding 1 in the head-on genes of each species (Table S2). Yet the total number of genes in each species was too small to establish statistical significance, necessitating a combined analysis of all data points. This analysis yielded a highly significant difference between the two groups, indicating that positive selection acts on a higher percentage of head-on genes (Table 1). Therefore, these data are consistent with positive selection driving the retention of at least some head-on alleles after spontaneous inversion from the opposing strand.

The cutoff value of dN/dS > 1 is a conservative criterion for identifying positive selection. The average dN/dS ratio is significantly higher for head-on genes across the genome regardless of whether we include the genes with a dN/dS>1. This difference remains even after the dN/dS >1 genes are removed from the data set, showing that the 2.4% of genes identified in Table 1 are not solely responsible for the higher dN/dS ratio of head-on genes (Table S3). Therefore, the dN/dS data provides strong evidence that (in addition to the 2.4% of head-on genes identified in Table 1) a higher percentage of head-on genes are under positive selection compared to co-directional genes.

**Table 1.** Positive selection acts on a higher percentage of head-on genes. The number of genes with a dN/dS ratio exceeding 1 were tallied across all six species and combined. Single tailed Chi-Squared test without a Yates correction was used to determine significance.

|  | Total Genes | # Genes<br>dN/dS >1 | Percent | Chi Sq. |
| --- | --- | --- | --- | --- |
| CD | 15,627 | 289 | 1.8 | 0.0039 |
| HO | 9,757 | 234 | 2.4 |  |

### Head-on genes are retained over evolutionary time scales

Given the detrimental effects of head-on transcription, it is possible that head-on genes may rapidly revert to the co-directional orientation. Previous work demonstrated that the negative GC skew of inverted DNA fragments rises due to normal replication, just like any other DNA fragment. Therefore, if the inverted regions identified here are retained for long time periods in the new orientation, their GC skew should have increased over time. Conversely, if inverted regions rapidly re-invert to the original orientation, their average GC skew value should mirror that of genes in the original orientation. Therefore, we compared the average GC skew of retained co-directional genes (those with a positive GC skew) to head-on genes that are the result of an inversion (those with a negative GC skew) (Fig. 5). These data show that the two groups do not have equivalent average GC skew values. Specifically, the head-on gene average is significantly higher (decreasing the magnitude) than predicted by the null hypothesis. This indicates that on average, head-on genes are retained for significant time periods following inversion. As such, these data are consistent with the possibility that head-on alleles can be beneficial.

### Head-on genes are enriched in common functions across species

To better understand the implications of co-directional to head-on gene inversions, we identified functions enriched among head-on genes across species (Fig. 6). To better evaluate consistency, we expanded the number of analyzed species (listed in the methods section). The DAVID bioinformatics database identifies a lower number of functional categories enriched among lagging strand genes: a ratio of 1:3 for head-on vs. co-directional genes, respectively (Fig. 6A). This trend is consistent between low and high co-orientation bias organisms which possess different overall ratios of head-on to co-directional genes. Gram-positive species possess roughly 1 head-on gene for every 5 co-directional genes, whereas the ratio is as low as 1:1.4 for low co-orientation bias organisms such as Gram-negative species. The consistency in the relative number of enriched functions between these diverse bacterial species suggests that gene function may be an important factor driving the segregation of genes between the leading and lagging strands.

**Figure 6.**
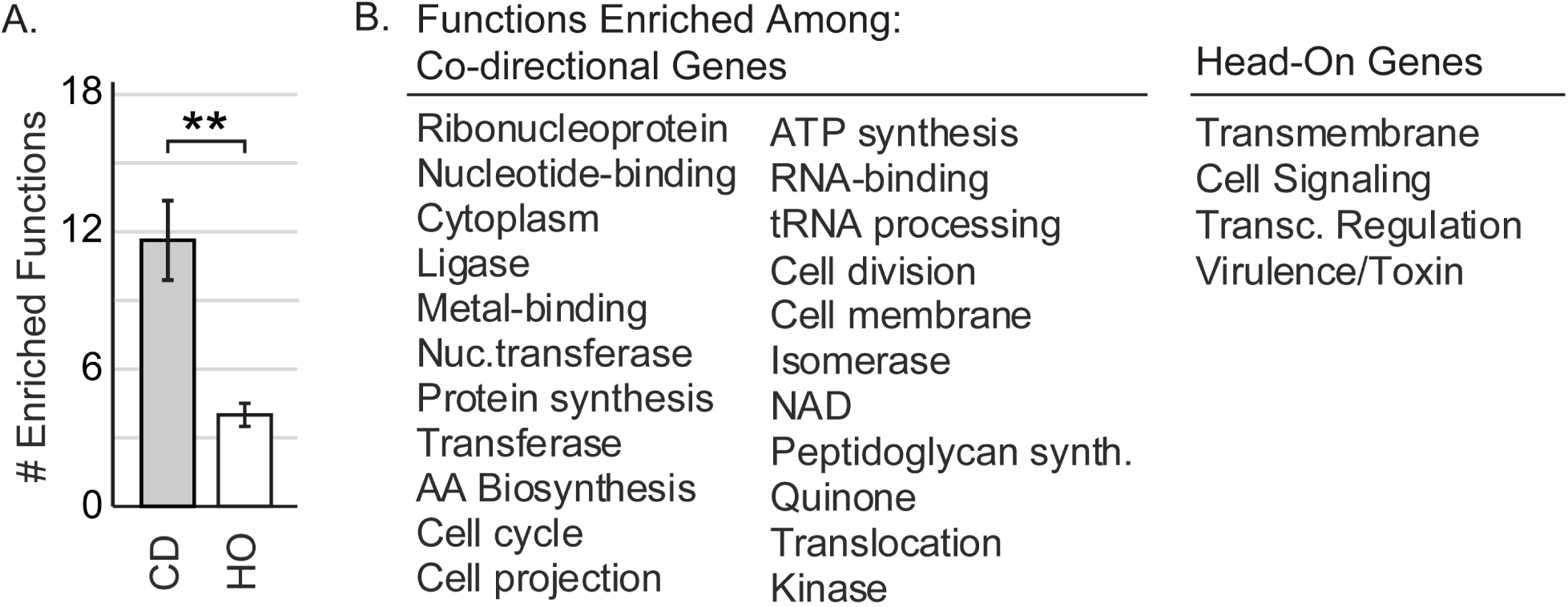
The functions enriched among head-on genes are consistent across species/phyla. A) The relative number functions enriched among co-directional (CD) versus head-on (HO) genes is highly consistent across phyla. The number of functions found to be enriched among CD or HO genes was determined for 12 species (species are listed in the methods section), then averaged across species. Error bars represent the standard error of the mean. Student’s t-test was used calculate statistical significance, ** indicates p-value = 0.0004. B) The functions of all lagging strand (head-on) or leading strand (co-directional) genes were determined on a per species basis for 12 bacterial species using the DAVID bioinformatics database (species are listed in the methods section). Functions found to be enriched in 4 or more organisms were included here as consensus functions.

We then found that the same functions are often enriched among head-on or co-directional genes between species (Fig 6B). Again, this suggests strongly that lagging strand encoding is non-random. Enriched functions include transcription regulation, trans-membrane localization, cell signaling, and virulence. Notably, though not overtly identified via the DAVID database analysis, we also find a considerable number of antibiotic resistance related genes including multidrug resistance pumps are encoded on the lagging strand (Table S4). This suggests that many virulence and antibiotic resistance genes may evolve at an accelerated rate, at least partially due to their head-on orientation.

### Many important virulence and antibiotic resistance genes are head-on

In order for replication-transcription conflicts to occur, both processes must happen simultaneously. Virulence genes are often transcriptionally induced during infection, a time when cells must also replicate their DNA. As such, we hypothesize that the expression of virulence genes during replication within the host should frequently cause replication-transcription collisions. In support of this premise, we previously showed that several stress response and virulence genes are head-on in *Listeria monocytogenes^13^.* Accordingly, we also found that the conflict resolution factor RNase HIII is essential for cell survival during infection of mice, and activated macrophages^13^. Here we provide additional evidence that a variety of antibiotic resistance and virulence genes anticipated to be expressed during infection are head-on (Table S4). Many of them are well characterized and are critical for virulence. Anecdotal examples include: *M. tuberculosis* multidrug resistance pump Rv1458c, which confers potent resistance to multiple front line antibiotics when overexpressed (dN/dS = 2.72, GC skew = -0.003)^37^; the *C. jejuni* siderophore transport protein *tonB3*, which is required for infection and has been retained in the head-on orientation longer than nearly all other genes in the genome according to its GC skew (GC skew = 0.23)^38^; the recently inverted distending toxin genes *cdtABC* (GC skew = -0.011)^39^; the S. *aureus* master virulence regulator *traP* (GC skew = 0.05) and *agrA* (GC skew = -0.15) which together regulate the small RNA virulence gene regulator, RNA III (co-directional)^40,41^. These examples serve to demonstrate anecdotally that important virulence genes are likely to be subject to head-on conflicts in nature. By extension, these data suggest that the head-on orientation of these genes can drive the rapid evolution of virulence and antibiotic resistance through head-on conflict-mediated increase in mutation rates.

## Discussion

Previous studies have demonstrated that most, if not all bacteria, avoid the worst head-on conflicts by co-orienting the transcription of highly expressed genes with replication. This co-orientation bias in essential genes should protect them from the high mutation rate conferred by head-on conflicts^4,19,42^. Yet the gene inversion patterns identified here indicate that the complete abolishment of head-on transcription is not advantageous. Instead, our findings highlight a surprising and fundamental pattern in the evolutionary history of bacterial species: the widespread creation of new head-on genes and operons via inversion. These observations are particularly interesting in light of a large body of work on replication-transcription conflicts which collectively set up the expectation that new head-on alleles should generally experience negative selection pressure due to their tendency to cause severe replication stress^1,2,13,42,43^. The data presented here directly contradict this expectation. As such, our work offers a major fundamental insight into the evolution of genomic architectures.

### Gene inversions and the evolution of genomic architecture

The findings presented here also raise important questions about the evolution of genome architectures and the existing co-orientation bias of genes in bacteria. For example, it is possible that the number of head-on genes is increasing relative to the number of co-directional genes. Alternatively, it is also possible that as new co-directional to head-on inversions occur, an equal number of head-on genes are inverting to the co-directional orientation. This could produce an orientation balance, ultimately maintaining the currently observed co-orientation bias of each species. However, the GC skew analyses alone, without extensive phylogenetic studies, cannot address the net change in the number of head-on to co-directional genes over evolutionary time.

### Possible second order selection for head-on alleles

We found that a higher percentage of head-on genes are under positive selection. For these genes, we propose that the elevated mutation rate conferred by head-on conflicts led to the more rapid production of advantageous mutations. This could have allowed positive selection to *indirectly* select for cells harboring the head-on allele^44,45^. It is important to re-iterate that we are not proposing that the head-on orientation is “used” to promote adaptation. Instead, the common creation/retention of head-on alleles is likely a product of second order selection, or the hitchhiker effect. This model offers a compelling explanation for the consistent retention of head-on genes in bacteria across phyla.

### Increasing evolvability through head-on replication-transcription conflicts

Evolvability, or evolutionary potential, can be defined as the relative ability of a particular strain to increase its fitness after evolving over a defined time period^44^. Experiments have demonstrated that in certain circumstances, increased mutation rates can increase evolvability, presumably through the more rapid creation of beneficial mutations (and despite the likely creation of more neutral/detrimental mutations)^44–48^. Based both on the data presented here, and previous work on conflicts, we propose that the continued inversion of genes to the head-on orientation increases the overall mutation rate and subsequently, evolvability of bacterial genomes. (Note that by “increase”, we are comparing existing cells which harbor numerous new head-on alleles to a hypothetical case in which the same strains retain the original corresponding co-directional alleles.)

### Potential mechanistic basis for gene inversions

As this study is particularly concerned with the inversion of whole genes and operons, we considered possible mechanisms through which the inversion of whole genes, rather than gene fragments, might occur. In particular, we looked for homology regions up/downstream of gene regions that could be used for recombination. We noticed that promoter and 3` UTR regions, by virtue of their AT richness, should have lower sequence complexity than regions in which all four nucleotides are equally distributed, i.e. within open reading frames. The depletion of guanine and cytosine in these regions should therefore increase the likelihood that upstream and downstream regions will be homologous. As such, the AT richness of promoters and 3` UTRs should be protective to open reading frames, promoting whole gene or operon inversion by homologous recombination, and discouraging truncations.

### Consistently enriched functions suggest that co-directional to head-on gene inversions can be adaptive

Interestingly, our functional analyses offer further support for the potential benefits of certain gene inversions. It appears that a specific subset of genes are particularly well suited for lagging strand (head-on) encoding: environmental sensing and stress response genes. These largely consist of trans-membrane proteins, transcriptional regulators and virulence genes. It is possible that these genes may be more mutable than other genes, that they rarely produce head-on conflicts, or that they have access to a higher number of mutations capable of providing a benefit. As head-on genes are a diverse group, all three possibilities may be correct for a subset of genes. Support for the adaptive hypothesis includes our identification of an increased frequency of positive selection acting on head-on genes and the enrichment of virulence genes on the lagging strand. The potential promotion of virulence gene mutations is consistent with the concept of the “everlasting” host-pathogen arms race^49–51^. This colloquial terminology describes the constant requirement for pathogens to rapidly create adaptive mutations in order to evade changing host immune responses. The concept of the evolutionary arms race and observed enrichment of virulence genes on the lagging strand together suggest that leading to lagging strand gene inversion may be a general mechanism capable of accelerating bacterial evolution and increasing the virulence of important pathogens.

## Methods

### Chromosome mapping

gView version 1.7 was used to map the GC skew and genes in Figure 1.

### Orthologue comparison

TimeZone v1.0 software was used to identify orthologues between 55 fully assembled *M. tuberculosis* genomes. Custom Python scripts were used mine strand and location information for each orthologue from corresponding Genbank files to identify inversions.

### Mutational Analysis

Ten or more fully assembled genomes were analyzed per species using TimeZone v1.0. Core genes were defined as having 95% similarity in amino acid content and gene length. Custom Python and Matlab scripts were used to analyze the location of *ori* and *ter* (inflection points in the chromosomal GC skew map) to assign gene orientation. Analyzed genomes are listed in Table S5.

### Functional enrichment analysis

The DAVID bioinformatics database was used to analyze functions enriched among all head-on or co-directional genes (not just core genes) of 12 species: *Mycoplasma gallisepticum* (we use *M. gallisepticum as a* proxy for the human pathogen *Mycoplasma genitalium.* Few genomes are available for *M. genitalium*, precluding a robust analysis), *Bacillus subtilis, Staphylococcus aureus, Listeria monocytogenes, Clostridiodes difficile* (formerly *Clostridium difficile), Pseudomonas aeruginosa, Klebsiella pneumoniae, Escherichia coli, Borrelia burgdorferi, Campylobacter jejuni, and Mycobacterium tuberculosis.* The output list of functions were manually edited to reduce the incidence of redundant functions. Specifically, we combined the following: membrane, transmembrane and membrane transport; transcription and transcription regulation; cell cycle and cell division; amino acid synthesis and the synthesis of any specific amino acid; metal binding and the binding of any specific metal ion.

## Acknowledgements

We would like to thank Sarah Mangiameli for her help with the *ori* and *ter* assignments. This work was supported by the Bill and Melinda Gates Foundation, Grant#OPP1154551, and National Institute of General Medical Sciences Award DP2GM110773 (to H.M.).

